# An automated model test system for systematic development and improvement of gene expression models

**DOI:** 10.1101/193367

**Authors:** Alexander C. Reis, Howard M. Salis

## Abstract

Gene expression models greatly accelerate the engineering of synthetic metabolic pathways and genetic circuits by predicting sequence-function relationships and reducing trial-and-error experimentation. However, developing models with more accurate predictions is a significant challenge, even though they are essential to engineering complex genetic systems. Here we present a model test system that combines advanced statistics, machine learning, and a database of 9862 characterized genetic systems to automatically quantify model accuracies, accept or reject mechanistic hypotheses, and identify areas for model improvement. We also introduce Model Capacity, a new information theoretic metric that enables correct model comparisons across datasets. We demonstrate the model test system by comparing six models of translation initiation rate, evaluating 100 mechanistic hypotheses, and uncovering new sequence determinants that control protein expression levels. We applied these results to develop a biophysical model of translation initiation rate with significant improvements in accuracy. Automated model test systems will dramatically accelerate the development of gene expression models, and thereby transition synthetic biology into a mature engineering discipline.

## INTRODUCTION

It has been a grand challenge to transition synthetic biology into a mature engineering discipline, where genetic systems (e.g. biosensors^1,2^, genetic circuits^3,4^, and metabolic pathways^5-7^) are reliably designed, built, and tested to reprogram cellular behavior with desired outcomes. Quantitative models play a central role in synthetic biology’s design-build-test cycle by predicting the function of a candidate genetic system, before it’s constructed, and therefore reduce trial-and-error experimentation. Gene expression models have been developed to predict the sequence-function relationships for several gene regulatory parts^2,8-11^, enabling the automated design of genetic systems with desired protein expression levels. Gene expression predictions have also been combined with system-level models of genetic circuits and metabolic pathways to predict how changes in system architecture, component expression levels, and host genome control organismal sensing, decision-making and the biosynthesis of desired chemicals^4,7,12-15^. Overall, more accurate gene expression models are becoming essential to correctly build genetic systems with many interacting components.

In mature engineering disciplines, automated test systems are routinely used to verify that models, design algorithms, and software systems generate predictions and outcomes within specified performance requirements^16^. Model test systems are run whenever an existing model is modified or when new models are proposed to ensure consistent improvements in accuracy across the widest possible range of inputs and to verify identical predictions across different software implementations. Model test systems also accelerate the discovery of new interactions by identifying the factors that contribute to model error. Within the life sciences, the CASP^17^, DREAM^18^, and IMPROVER^19^ contests and competitions have served a somewhat analogous purpose, where researchers are challenged to apply computational modeling to solve complex problems, for example, predicting protein structure from sequence, identifying disease genetic traits, and reverse-engineering gene regulatory networks. A key theme of these contests is that truly novel mechanisms are far more discoverable once state-of-the-art models are challenged to predict the outcome of a large and diverse experimental dataset.

Here, we present the first automated test system for gene expression models and apply it on a compiled database of 9862 characterized genetic systems with highly diverse DNA sequences and measured functions in diverse host organisms. Beyond statistical analysis, we use information theory to critically assess the amount of information added by a model’s predictions, automated model testing to accept or reject a large set of mechanistic hypotheses, and machine learning to identify new sequence determinants and mechanisms. We demonstrate these capabilities on six sequence-to-function models of bacterial translation initiation^20-25^, providing the first systematic test of their accuracies, and explaining the mechanistic origins of the models’ error. Ultimately, we identified several mechanisms missing from the state-of-the-art model, which led to the development of a novel model of bacterial translation initiation with significant improvements in accuracy. Overall, we show that the model test system accelerates the systematic development of improved gene expression models.

### An automated model test system for gene expression models

We first compiled a database of 9862 characterized genetic systems, collected from 15 publications^2,21,23-35^, whereby 7 unique heterologous reporter proteins (mRFP1, super folder GFP, CFP, YFP, LacZ, Luciferase, and NanoLuc) were expressed using engineered promoters and ribosome binding sites in 6 different bacterial hosts (*E. coli*, *B. subtilis* 168, *C. glutamicum*, *S. typhimurium* LT2, *B. thetaiotaomicron*, and *P. fluorescens*). The genetic systems’ reporter expression levels were individually characterized using flow cytometry or spectrophotometry, or using FlowSeq, a technique that uses binned cell sorting, bar-coding, and next-generation sequencing to measure the transcription and translation rates of a library of genetic systems^29^. For FlowSeq measurements, we applied automatic filters to remove measurements with insufficient or skewed read counts (**Methods**). We then grouped together genetic systems first by their characterization method and second by their commonalities (same promoter, same reporter protein, and same host) to create two high-quality databases: 1014 individually characterized genetic systems (1014IC) with 22 sub-groups and 8848 FlowSeq characterized genetic systems (8848FS) from a single experiment (**Supplementary Table 1**).

We then developed an automated model test system that reads each genetic system database and uses its sequence information to predict the genetic systems’ protein expression levels, using one of many selectable drop-in quantitative gene expression models (Figure 1). These gene expression models were developed with the expectation of a linear relationship between the predicted and measured expression levels, though the relationship’s proportionality constant will vary by sub-group, for example, due to the utilization of different reporter proteins in different host organisms. We therefore developed an approach to automatically identify the correct proportionality constant for each sub-group, utilizing median absolute deviation (MAD) criteria to reject outliers and five common statistical metrics (Table 1) to quantify each model’s overall accuracy and precision (**Methods**). Motivated by our observations below, we then augmented the model test system to perform a more advanced information theoretic analysis that additionally accounts for the diversity of the genetic systems’ sequences and the precision of the experimental measurement techniques. By calculating both commonly used and newly proposed metrics for model accuracy, the model test system enables researchers to view each metric with the necessary context to make appropriate comparisons.

**Table 1:** The model test system accuracy and precision metrics for seven models of bacterial translation initiation rate, evaluated on the 1014IC and 8848FS datasets. The range for R^2^ and the normalized Kullback-Leibler divergence (*D*_*KL*_) is (0,1). The range for AUROC is (0.5,1). The root mean square error (RMSE) is always positive. The model’s one-sided error cumulative distribution function is provided at selected thresholds. NP is the number of genetic systems where the model is not capable of calculating a prediction.

| 1014 Individually characterized (1014IC) |  |  |  |  |  | % within X-fold error |  |  | NP |
| --- | --- | --- | --- | --- | --- | --- | --- | --- | --- |
| Model | $R^2$ | RMSE | $D_{KL}$ | AUROC | MC (bits) | 2 | 5 | 10 | |
| RBS Calculator v1.0 | 0.51 | 1.75 | 0.41 | 0.85 | 3707 | 35 | 66 | 79 | 66 |
| RBS Calculator v1.1 | 0.66 | 1.51 | 0.46 | 0.89 | 5911 | 44 | 77 | 88 | 23 |
| RBS Calculator v2.0 | 0.71 | 1.39 | 0.48 | 0.93 | 7313 | 50 | 81 | 90 | 23 |
| RBS Calculator v2.1 | 0.74 | 1.31 | 0.49 | 0.93 | 7857 | 49 | 82 | 91 | 0 |
| UTR Designer | 0.50 | 1.77 | 0.43 | 0.85 | 4490 | 38 | 69 | 80 | 66 |
| RBS Designer | 0.42 | 1.90 | 0.35 | 0.85 | 2385 | 33 | 57 | 69 | 101 |
| EMOPEC | 0.31 | 2.14 | 0.40 | 0.76 | 3053 | 31 | 64 | 77 | 1 |
| 8848 FlowSeq characterized (8848FS) |  |  |  |  |  | % within X-fold error |  |  | NP |
| Model | $R^2$ | RMSE | $D_{KL}$ | AUROC | MC (bits) | 2 | 5 | 10 | |
| RBS Calculator v1.0 | 0.25 | 1.03 | 0.21 | 0.80 | 1851 | 39 | 74 | 88 | 18 |
| RBS Calculator v1.1 | 0.26 | 1.03 | 0.22 | 0.80 | 1822 | 40 | 75 | 88 | 0 |
| RBS Calculator v2.0 | 0.26 | 1.02 | 0.22 | 0.78 | 1421 | 36 | 71 | 86 | 0 |
| RBS Calculator v2.1 | 0.28 | 1.01 | 0.22 | 0.78 | 1982 | 40 | 78 | 92 | 0 |
| UTR Designer | 0.23 | 1.04 | 0.21 | 0.80 | 2082 | 42 | 79 | 91 | 19 |
| RBS Designer | 0.23 | 1.05 | 0.27 | 0.73 | 511 | 23 | 52 | 70 | 0 |
| EMOPEC | 0.09 | 1.14 | 0.21 | 0.64 | 2782 | 47 | 86 | 95 | 1 |

**Figure 1.**
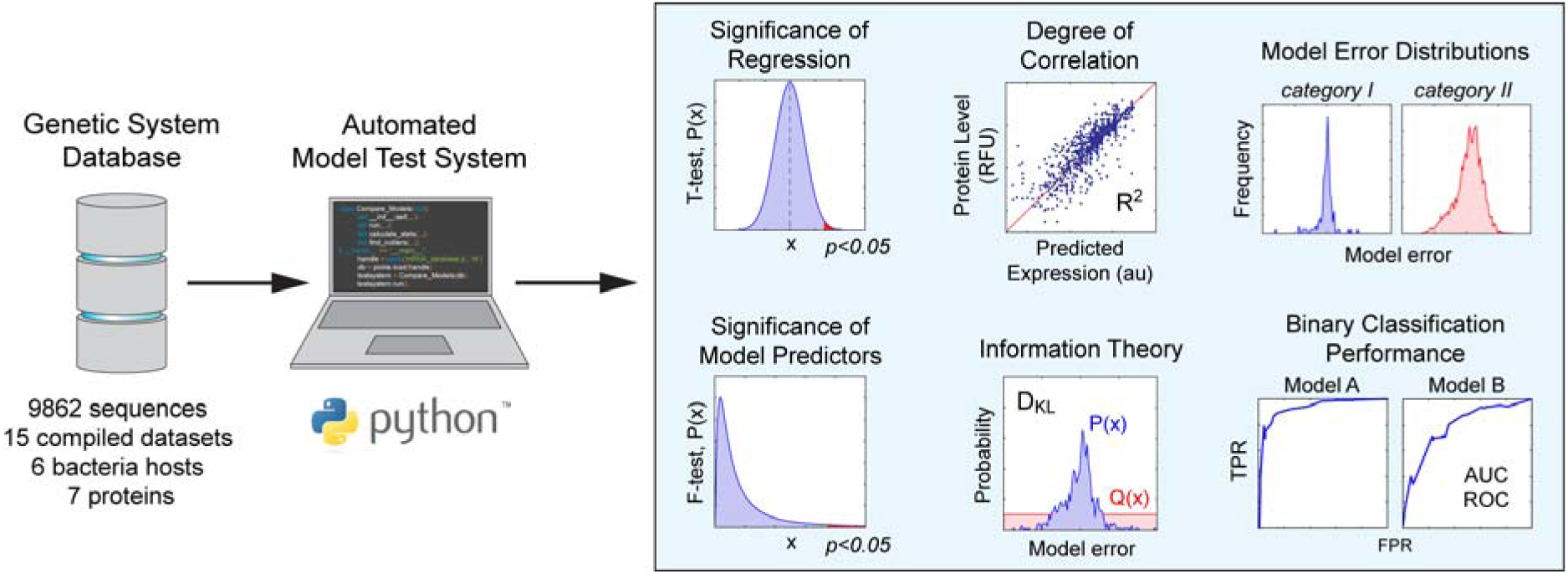
The inputs and outputs of the model test system. The database contains characterized genetic systems and relevant metadata, including their genetic part sequences, host organism specifications, experimental conditions, and publication source in machine- and human-readable formats. The model test system systematically evaluates the predictions of proposed models and returns each model’s accuracy statistics, error distributions, and information theory metrics. These calculations are then automatically utilized to evaluate hypotheses, categorize sequences, and identify sources of model error.

### A systematic comparison of translation initiation rate models

Several known ribosome-mRNA interactions work together to control an mRNA’s translation initiation rate, including preliminary binding of the 30S ribosomal subunit to upstream standby sites in the mRNA (interaction #1)^24,36^, unfolding of inhibitory mRNA structures (interaction #2)^37^, hybridization between the ribosome’s 16S rRNA and the mRNA at the Shine-Dalgarno sequence (interaction #3)^38^, hybridization between the tRNA^fMet^ and start codon (interaction #4)^39^, and entropic stretching or compression of the ribosome caused by non-optimal distances between the Shine-Dalgarno and start codon (interaction #5)^40^. However, there is considerable debate over the importance of these interactions, the best way to calculate their strengths, and whether there are additional interactions that control translation rate. Thus, several models of translation initiation have been proposed with both conceptual and implementation differences.

Using the automated model test system, we evaluated the accuracies and sources of error for six different models of bacterial translation initiation rate^20-25^ on both the 1014IC and the 8848FS datasets. The RBS Designer^22^ uses dynamical systems theory to model ribosome binding, calculating the ribosome binding site’s exposure probability (interaction #2) and the free energy of mRNA-rRNA hybridization (interaction #3) to predict an mRNA’s translation initiation rate. The RBS Calculator (v1.0) uses statistical thermodynamics and a free energy model to calculate the ribosome’s total binding free energy to an mRNA sequence, and predict the mRNA’s translation initiation rate, accounting for interactions #1 to #5. The RBS Calculator v1.1^23^ uses a modified free energy model implementation that includes the unfolding of inhibitory mRNA structures within the protein’s coding sequence (interaction #2) and a new approach for calculating hybridization free energies that becomes important when mRNAs have non-canonical Shine-Dalgarno sequences (interaction #3). The RBS Calculator v2.0^24^ incorporates new biophysical rules governing the ribosome’s interactions at mRNA standby sites, determined through characterization of rationally designed mRNAs (interaction #1). The UTR Designer^20^ is a modified version of the RBS Calculator v1.0 that introduces a direct and indirect pathway to unfolding inhibitory mRNA structures (interaction #2). EMOPEC^25^ is an empirical model that uses a look-up sequence table of measured hybridization free energies (interaction #3) and the model of non-optimal spacing from RBS Calculator v1.0 (interaction #5) to predict an mRNA’s translation initiation rate. Detailed model descriptions can be found in **Supplementary Discussion 1**.

The model test system revealed clear differences in model accuracies and highlighted the utility of applying multiple test metrics to evaluate model performance (Table 1). When evaluated on the 1014IC dataset, the successive modifications to the RBS Calculator model resulted in accuracy improvements that were reflected across all test metrics. To compare, the test metrics largely agreed that the UTR Designer, RBS Designer, and EMOPEC models have lower accuracies (Figure 2a). Overall, the RBS Calculator v2.0 model could predict 50% of the 1014IC mRNAs’ translation initiation rates within 2-fold and 90% within 10-fold with extremely high classification accuracy, an AUROC of 0.93 (**Supplementary Figure 1**). Notably, the RBS Calculator v2.0 had variable accuracies on data sub-groups utilizing different reporters, measurement techniques, and host organisms, motivating the further development of automated testing strategies to learn why these discrepancies exist (**Supplementary Figure 2**).

**Figure 2.**
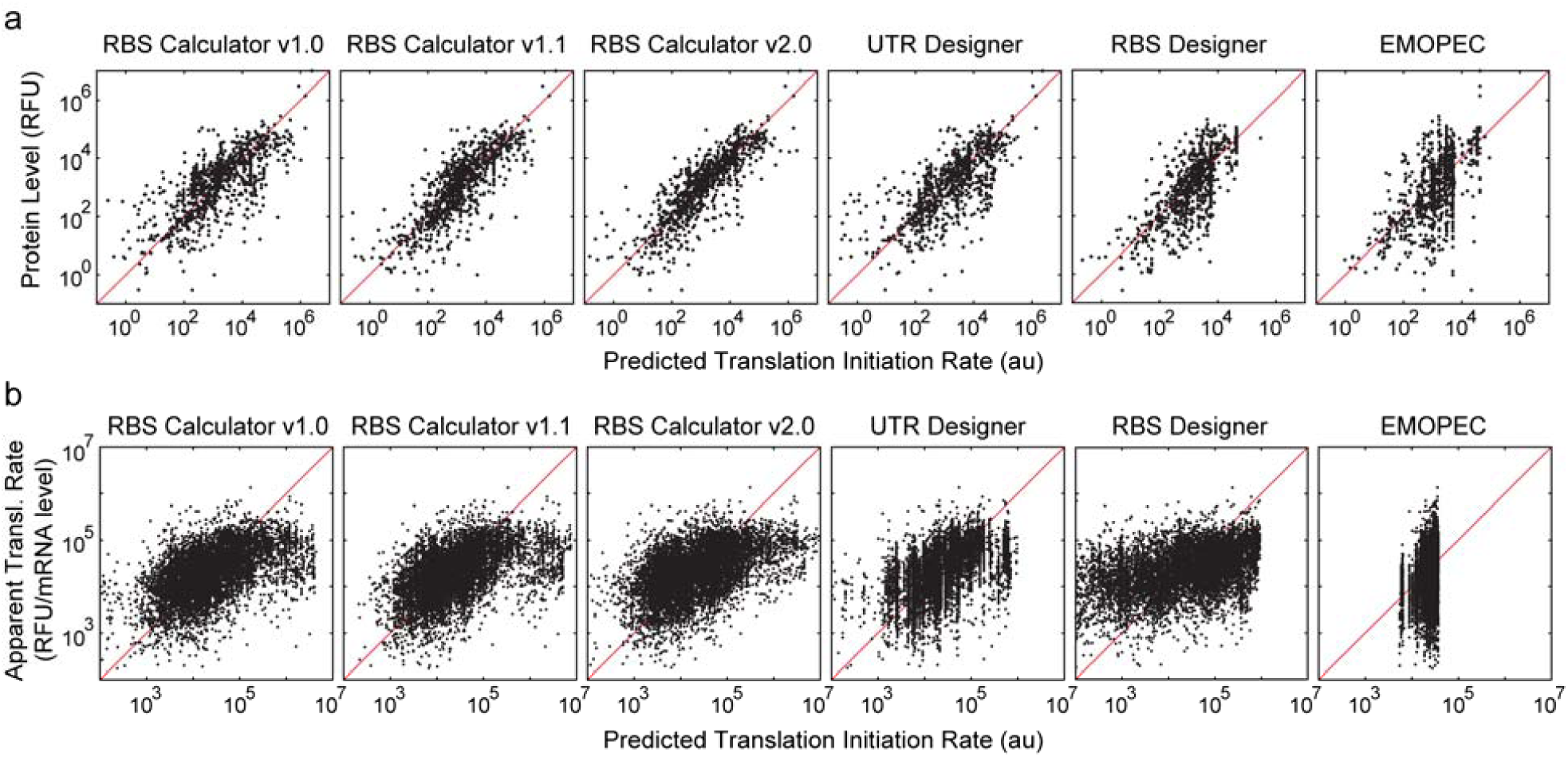
Systematic assessment of six sequence-to-function models of bacterial translation initiation rate evaluated on the (**a**) 1014IC and (**b**) 8848FS datasets. Test metrics are listed in Table 1.

However, when evaluated on the 8848FS dataset, there was an overall disagreement among the test metrics over the accuracies of each model (Figure 2b, Table 1). The Pearson R^2^, D_KL_, and AUROC metrics show that all six models had similarly sharp reductions in accuracy with only slight differences between them, compared to lower RMSEs indicative of higher accuracy. Notably, these metrics also had a poor correspondence to the models’ error distributions. In particular, while EMOPEC’s test metrics showed the lowest model accuracy, its model error distribution had the highest percentage of well-predicted sequences, clearly contradicted by a comparative graphical analysis (Figure 2b). Overall, these results show that once experimental datasets grow beyond a certain size, current approaches to quantifying a model’s accuracy strongly depend on the chosen test metric, creating a subjective choice in what should be an objective process. All dataset sequences, measurements, and model calculations are listed in **Supplementary Data 1** and **Supplementary Data 2**.

### Information theory to determine model capacity

Using smaller scale examples, we first investigated how the same model evaluated on different data sub-groups had substantially different apparent accuracies. For example, the EMOPEC model could accurately predict the gene expression levels of a 10-sequence *E. coli*-LacZ sub-group (R^2^ = 0.85) where mutation of only 3 nucleotide positions resulted in a 2400-fold change in translation rate. Similarly, the RBS Calculator v2.0 model was extremely accurate when evaluated on a 14-sequence *P. fluorescens*-mRFP1 sub-group (R^2^ = 0.90) where mutation of 4 nucleotide positions resulted in a 10900-fold change in translation rate (**Supplementary Figure 3**). However, even though these models tested well on datasets with a small number of nucleotide mutations, they did not achieve similar accuracies on larger, more diverse datasets (**Table I**). These results inspired us to measure the difficulty of correctly predicting the expression levels of a genetic system dataset. From an information theoretic perspective, a model’s accuracy is determined by its ability to maximally reduce uncertainty in the predicted outcomes. It is therefore equally important to consider both the starting amount of uncertainty in the dataset and the ending amount of uncertainty in the gene expression level predictions.

We therefore derived a new test metric, called Model Capacity (*MC*), which quantifies a model’s ability to accurately predict gene expression levels by tracking the flow of information from source to comparator and measuring the model’s ability to reduce uncertainty in the predicted outcomes, taking into account the genetic system dataset’s sequence diversity, the experimental measurements’ functional diversity, and the model’s error distribution (**Figure 3a**). To calculate *MC*, we quantify a dataset’s sequence diversity (*H*_seq_) utilizing Shannon’s entropy, we determine a dataset’s number of distinguishable outcomes (*N*) by dividing the measurements’ dynamic range by its average precision, and we compute the Shannon entropy of the model’s error distribution, compared to a random model, both using *N* outcomes (see **Supplementary Discussion 2** for a detailed explanation). Overall, our newly proposed *MC* metric provides an objective way to assess both the model and the dataset together, facilitating correct cross-data model comparisons.

For example, in agreement with currently used test metrics, the successive improvements to the RBS Calculator model led to higher MCs of 3707, 5911, and 7313 bits when evaluated on 1014IC. The MCs for the UTR Designer, EMOPEC, and RBS Designer models also closely tracked the other test metrics (4490,3053, and 2385 bits, respectively) on this dataset (Figure 3b). However, only the MC metric can assess whether a dataset has sufficient sequence and functional diversity to truly test a model’s accuracy. For example, both the RBS Calculator v2.0 and EMOPEC models have very low MCs when evaluated on the 14-sequence *P. fluorescens*-mRFP1 data sub-group (33 vs. 31 bits, respectively). Similarly, the RBS Calculator v2.0 and EMOPEC models have even lower MCs when evaluated on the 10-sequence *E. coli*-LacZ data sub-group (14 vs. 9 bits, respectively). The MC metric correctly assessed that these small datasets are too homogenous to provide a distinguishable comparison in model accuracy; the relative differences in MC are too small, compared to the absolute MC values possible when evaluated on the more diverse 1014IC dataset.

**Figure 3.**
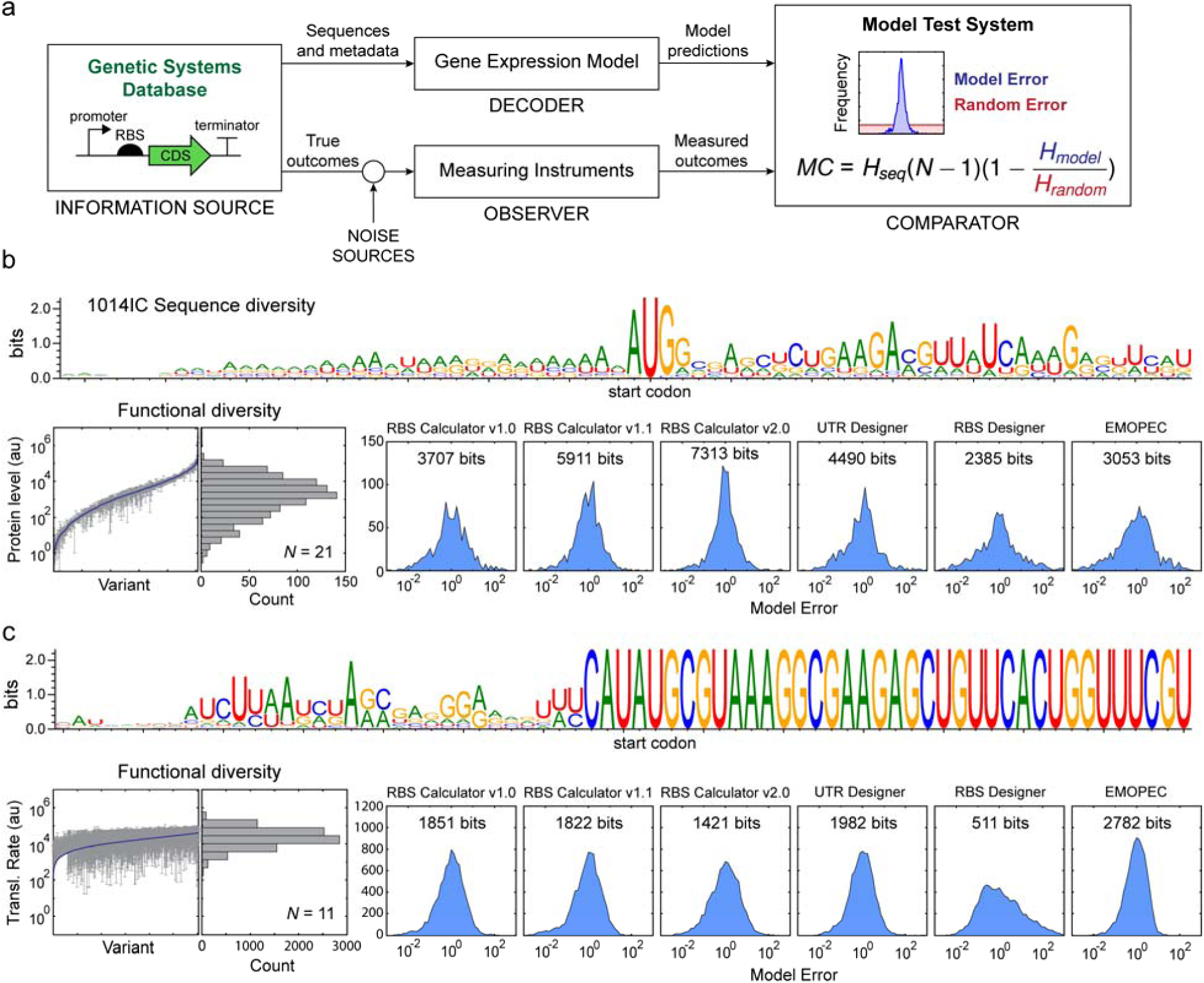
Information theory to quantify dataset uncertainty and a model’s capacity (*MC*) to predict outcomes.(**a**) A schematic showing the flow of information from a list of genetic systems (the source) to the model test system (the comparator), using digital systems to predict outcomes (the decoder) and the physical systems to measure outcomes (the observer). *N* is the number of distinguishable measurement outcomes. *MC* is the reduction in uncertainty from information source to model error comparator. The sequence logos^41^, functional diversities, error distributions, and model capacities for (**b**) the 1014IC and (**c**) the 8848FS dataset.

Notably, a dataset with more sequences does not always lead to higher sequence diversity. For example, the smaller 1014IC dataset has a larger sequence diversity (H_seq_ = 2243 bits), compared to 8848FS (H_seq_ = 1448 bits), because it contains longer 5’ UTRs with mutations at more distributed positions (Figure 3b-c). Moreover, different characterization techniques can also have substantially different dynamic ranges and measurement precisions. For example, while the FlowSeq technique simultaneously measures gene expression levels from thousands of sequences, the current approach sorts fluorescent cells into a relatively small number of defined bins, resulting in a smaller number of distinguishable outcomes, compared to measuring reporter expression levels within isogenic cultures (*N* = 11 for FS8848 vs. *N* = 21 for 1014IC). Together, *H*_*SEQ*_ and *N* measure the maximum possible uncertainty in a dataset, and therefore control the maximum possible *MC* for a perfect model evaluated on that dataset (MC_max_ of 47100 bits for 1014IC, and 14476 bits for 8848FS). As a result, even the most accurate model (*MC*), when evaluated on 8848FS (EMOPEC) has a lower *MC*, compared to its evaluation on 1014IC (2782 vs 3053 bits respectively), a symptom of its limited sequence and functional diversity. Future efforts to test model predictions should therefore seek to design and characterize datasets with higher MC_max_.

### Automated hypothesis testing to evaluate mechanisms and model implementations

Beyond assessments of model accuracies, we next expanded our model test system to automatically apply statistical analysis to accept or reject proposed mechanistic hypotheses. Here, we demonstrate that capability by evaluating 100 mechanistic hypotheses covering the wide-ranging debate over the sequence determinants that control translation initiation rate. Each hypothesis was first automatically converted into a distinct model implementation, followed by evaluating its predictions on the 1014IC dataset. Hypotheses were accepted only when an *F*-test identified that the model implementation’s error distribution had a statistically significant reduction in variance (*F*-test *p* < 0.01) and when its accuracy test metrics were improved (higher R^2^ and MC), compared a RBS Calculator v2.0 baseline model. The conclusions for all 100 hypotheses are listed in **Supplementary Data 3**.

Surprisingly, when proposed as hypotheses, several commonly accepted anecdotes were rejected. It has been generally accepted that the ribosome binding site, and specifically, the Shine-Dalgarno sequence, is 6 nucleotides long, suggesting that only these nucleotides hybridize with the 3’ end of the 16S ribosomal RNA (SD: 5’-AGGAGG-3’, interaction #3). However, by constructing model implementations with 55 different possible 16S rRNA sequences, covering all possible Shine-Dalgarno sequences from 4 to 13 nucleotides long, our analysis revealed that the *E. coli* Shine-Dalgarno sequence is 9 nucleotides long (SD: 5’-UAAGGAGGU-3’, R^2^ = 0.70, *F* = 0.85, *p* = 0.013, **Supplementary Figure 4**). Several studies have also claimed that only the Shine-Dalgarno interaction or only the presence of mRNA structures are responsible for controlling a mRNA’s translation rate. Instead, a recursive feature reduction method showed that both interactions must be considered to accurately predict translation rates (**Supplementary Table 2**). A model that only incorporates Shine-Dalgarno interactions does not accurately predict translation rates (R^2^ = 0.54, *F* = 3.3, *p* = 3.6 × 10^−75^). Likewise, a model that only quantifies the energy needed to unfold mRNA structures is not accurate either (R^2^ = 0.44, *F* = 6.2, *p* = 4.6 × 10^−159^). Here, the model test system demonstrated its ability to strongly reject false claims with extremely high statistical significance.

**Figure 4.**
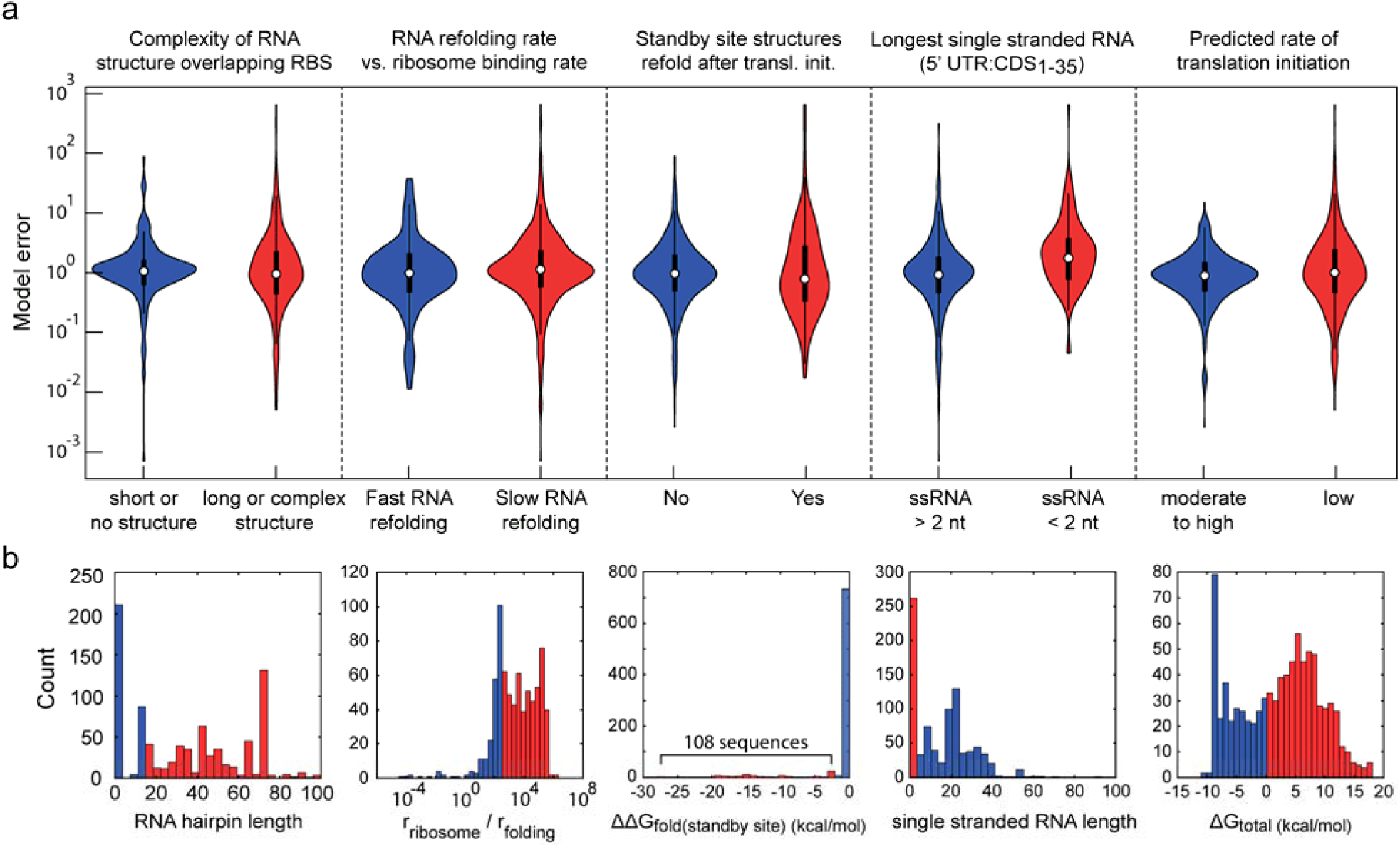
Identifying the sources of model error. (**a**) Violin plots comparing the error distributions from RBS Calculator v2.0 model predictions across five selected categories of mRNA sequences as shown. (**b**) Histograms illustrating the thresholds and counts of sequences for each of the categories as shown in (**a**).

Separately, translation rate model predictions have relied on the accurate calculation of RNA folding free energies and structures, though it is not known how these calculations affect the translation model’s accuracy. Several options are available; translation rate models may use different RNA free energy parameters, they may calculate either a minimum-free-energy (mfe) or ensemble centroid RNA structure, and they may incorporate additional energetic contributions, such as dangling nucleotide interactions. We therefore formulated and evaluated all combinations of these options as hypotheses. We found that utilizing the ensemble centroid RNA structure (R^2^ = 0.58, *F* = 4.7, *p* < 1.8 × 10^−119^) or incorporating the dangling nucleotide interaction (R^2^ = 0.69, *F* = 1.2, *p* = 0.002) resulted in less accurate predictions. We also found that the most recently developed RNA free energy parameter set^42^ provided a statistically significant improvement in model accuracy (R^2^ = 0.72, *F* = 1.2, *p* = 0.008).

Importantly, the model test system also determines when a proposed hypothesis is not testable on a given dataset.If a proposed mechanism or interaction is predicted to have the same, or similar, effect on all predicted outcomes within a data sub-group, then accepting or rejecting the hypothesis is not possible. We found two interesting examples of such hypotheses that could motivate future efforts. First proposed in 1990, the downstream-box (DB) hypothesis suggests that a portion of an mRNA’s CDS can hybridize to the 16S rRNA_1469:1483_ in *E. coli* to accelerate its translation initiation rate^43^. This hypothesis was untestable was because all data sub-groups share similar CDS regions, and therefore no matter how the interaction is proposed to take place, the predicted translation initiation rates within each data sub-group are similarly scaled. Second, it has been proposed that some organisms, including *Bacteroides thetaiotaomicron*, do not utilize a Shine-Dalgarno sequence to initiate translation^28,44^. We therefore evaluated the importance of the Shine-Dalgarno (interaction #3) when evaluated on the 143 mRNAs characterized in *B. thetaiotaomicron* and found that model accuracies were nearly identical with and without this interaction (R^2^ = 0.51 vs. 0.49, *F* = 1.05, *p* = 0.39).Here, the 143 mRNAs do not have sufficient sequence and functional diversity to test this hypothesis; specifically, the maximum change in interaction #3’s energy was relatively small, compared to the overall 1014IC dataset (4.2 vs. 18.6 kcal/mol).Overall, this capability of the model test system can accelerate mechanistic discoveries by directing the design and characterization of datasets to test previously untestable hypotheses.

### Sequence-function-error associations to identify new gene expression determinants

Once a gene expression model encompasses all well-characterized interactions controlling gene expression, it can become a significant challenge to discover and describe novel gene regulatory mechanisms that control gene expression levels. By combining the model test system with machine learning techniques, we next demonstrated how to systematically discover poorly characterized interactions controlling gene expression levels within the 1014IC dataset. We first enumerated a long list of candidate biophysical, biochemical, and geometric properties that can all be evaluated from sequence information and have the potential to influence gene expression. Our list included the characteristics of calculated structures of portions of mRNA, including the lengths, nucleotide compositions, and folding free energies of key structural features (hairpin loops, bulges, mismatches, single-stranded regions); the kinetic folding rates of these RNA structures as determined by the algorithm KFOLD^45^, and calculated characteristics of the ribosome-mRNA interaction, such as the structures of upstream standby sites^24^ and the compositions of the spacer region^34^ (**Supplementary Table 3**). The model test system then uses a feature selection method to automatically iterate over all candidate properties to identify ones where changes in the sequence-function relationship are associated with a statistically significant increase in the gene expression model’s error, here using the RBS Calculator v2.0 as the baseline. Overall, we identified five properties associated with increased model error that have sufficient statistical significance to warrant further investigation (*p* < 0.05) (Figure 4, **Supplementary Table 4**).

First, we found that long mRNA structures (>29 nucleotides long) that overlapped with the Shine-Dalgarno sequence greatly reduced the model’s ability to correctly predict the mRNA’s translation rate (*F* = 43.0, *p* = 2.8 × 10^−210^). These structures are inhibitory because they must be unfolded before the ribosome can initiate translation, and therefore any miscalculation in their unfolding free energies will have a large effect on the translation rate prediction. Relatedly, we found that whenever inhibitory mRNA structures required a large amount of time to fold (>322 KFOLD time units), or when the ribosome’s binding rate was faster than the RNA structure’s refolding rate, the model’s predictions were significantly less accurate (*F* = 12.3, *p* =5.5 × 10^−97^; and *F* = 40.1, p = 3.2 × 10^−115^ respectively). We also found that when standby site structures were predicted to refold to a lower energetic state following ribosome binding, then the model’s predictive accuracy was significantly reduced (*F* = 126.6, *p* = 8.4 × 10^−79^). During the cycling of translation initiation, it is possible that these structures do not have sufficient time to refold to the minimum free energy structure, or are kinetically trapped.

Finally, we found that highly structured 5’ untranslated regions with fewer than 2 consecutive single-stranded RNA nucleotides had less accurately predicted translation rates (F= 22.5, p = 5.0 × 10^−181^). Single-stranded RNA regions are potential binding sites for RNase E, and therefore the absence of such sites could increase mRNA stability and affect gene expression levels^46^. Similarly, we found that whenever mRNAs had low predicted translation initiation rates, then the model also had higher error (*F* = 450, *p* < 10^−100^), in agreement with evidence that poorly translated mRNAs become unprotected by ribosomes and therefore are more likely to be degraded by RNase E^47^. Altogether, the model test system identified 671 mRNAs (66% of the 1014IC dataset) where the changes in mRNA degradation and stability have confounded the translation rate model’s predictions, explaining a key source of error.

### The Model Test System Enables Systematic Development of Improved Models

We then leveraged the compiled genetic system database, confirmed mechanistic hypotheses, and identified sources of model error to develop and validate a new version of the RBS Calculator free energy model (v2.1) with a significant improvement in accuracy. The model test system enabled us to rapidly propose and evaluate new formulas for quantifying interaction energies that control ribosome binding and translation initiation rate, particularly new interactions that only became apparent when evaluated on a large genetic system database. Specifically, we added a new free energy term (ΔG_stack_) to quantify stacking interactions between adjacent RNA nucleotides, which had a large effect on ribosome binding sites with homopolymer sequences^34^ (**Supplementary Figure 5**); we improved the formulas that quantify how ribosomes in gram-positive organisms bind to structured standby sites and stretch/compress with varying ribosome binding site spacer lengths (**Supplementary Figure 6**)^48^; we improved the calculation of apparent tRNA^fMet^ binding free energies at non-canonical start codons, using 35 recently characterized mRNAs^35^ (**Supplementary Figure 7**) and we updated the translation rate model’s RNA free energy parameters to the Andronescu2007 set^42^. Importantly, while these improvements relied on characterization of mRNA translation rates using different plasmid or genomic copy numbers, promoters, reporters, and measurement techniques, the model test system could correctly extract the maximum amount of information from the measurements to parameterize and validate model predictions. The resulting RBS Calculator v2.1 model predictions were more accurate than v2.0 across the entire 1014IC dataset (R^2^ = 0.74 vs. 0.71, MC = 7857 vs. 7312 bits), including a statistically significant reduction in the model error distribution’s variance (F = 0.74, p = 9.8 × 10^−7^), with predictions for all sequences compared to 23 not predicted with v2.0 (**Supplementary Figure 8**).

Beyond these improvements, the model test system identified additional confounding factors, such as changes in mRNA folding kinetics and mRNA degradation rates, that are not considered in an equilibrium thermodynamic model of translation initiation rate, but nonetheless increase its model error. To avoid these sources of error, we then utilized the model test system’s categorization of sequence-error associations (Figure 4) to propose a safe operating zone where RBS sequences could be more accurately designed using the model. Using the mRNA sequence alone, a model prediction is considered outside the safe operating zone when the mRNA contains a long or slow-folding RNA structure (>29 nucleotides long or >320 KFOLD time units), when standby site structures are predicted to refold (ΔΔG_fold,standby site_ > 0), when mRNAs contain long single-stranded RNA regions (>5 nucleotides), and when the predicted translation initiation rate is low, resulting in greater deprotection of mRNAs (ΔG_total_ > 0). Altogether, the RBS Calculator v2.1 model’s accuracy was significantly improved when restricting its predictions to sequences within the safe operating zone. The model predicted the translation rates of 64%, 91% and 95% mRNAs to within 2-fold, 5-fold, and 10-fold of measurements, respectively, and with a statistically significant reduction in the model error distribution’s variance (F = 0.47, p = 9.9 × 10^−12^).

## DISCUSSION

The field of Synthetic Biology has considerably matured since its origins, though several cycles of designing, building, and testing organisms are still required to engineer complex genetic systems^14^. Gene expression models have greatly reduced the number of engineering cycles by designing sequences to control protein expression levels, and thereby rationally direct genetic system optimization^23^. However, more accurate gene expression models are needed to engineer complex genetic systems with many interacting components. For several reasons, it has been a significant challenge to systematically develop more accurate gene expression models. First, genetic systems are often characterized using different promoters, reporters, host organisms, and techniques, resulting in disparate datasets that require careful analysis to extract information. Second, several interactions collectively control sequence-expression relationships; correctly quantifying their strengths is a multi-variable, high dimensional problem. Finally, a model’s apparent accuracy depends on the dataset used to evaluate it, and statistical test metrics often disagree on accuracy improvements.

To overcome these challenges, we created an automated model test system that extracts the maximum amount of information from large genetic system databases, systematically assesses gene expression model accuracies, evaluates the validity of mechanistic hypotheses testing the importance of specific interactions, and identifies sequence-dependent physical properties that control a model’s error (Figure 1). We applied the model test system to carry out the first systematic assessment of six gene expression models that predict the translation initiation rates of mRNAs, showing that models have clear differences in accuracy across five common test metrics and two large datasets (Figure 2, Table 1).To explain these differences, we utilized information theory to quantify the uncertainty in both the dataset and the model predictions together, resulting in the first test metric, called Model Capacity (MC), that is capable of correct cross-dataset comparisons (**Figure 3**). Using advanced statistics and machine learning techniques, we then evaluated the validity of 100 mechanistic hypotheses and identified five sources of model error to distinguish model implementations and identify areas for model improvements (Figure 4). Using this analysis, we then rapidly developed a new version of RBS Calculator (v2.1) that incorporates several newly quantified interactions to yield statistically significant improvements in accuracy. These improvements in model accuracy considerably reduce trial-and-error experimentation, particularly when engineering complex genetic systems. For example, when using RBS Calculator v2.1 to engineer a 30-protein system with a 5-fold targeted expression operating space, only 2 design trials per protein are needed to yield a 78% chance of first-cycle success, compared to 5% when using RBS Calculator v2.0. Similarly, the success rate is increased to 98% when using 3 design trials per protein, compared to 41% (**Supplementary Discussion 4**). Overall, the model test system enables the accelerated analysis of model predictions on large datasets to focus model development on poorly characterized interactions that contribute most to model error.

The model test system is open-source software (<u>https://github.com/reisalex/SynBioMTS</u>), developed with the explicit goal of making it easy to expand the genetic system database, calculate additional test metrics, assess new gene expression models, evaluate additional hypotheses, and identify new sequence-dependent properties that correlate with model error. We also created an Excel template (**Supplementary Data 4**) to enable researchers to add characterized genetic systems to the growing database. Expanded genetic system databases will be made available periodically, which we envision will be used to develop and improve predictive sequence-to-function models for many additional gene regulatory mechanisms, including transcriptional regulation, termination, mRNA decay, gene expression coupling, and post-transcriptional regulation by small RNAs, riboswitches, and ribozymes.

## ACKNOWLEDGEMENTS

We would like to thank Daniel Goodman (Harvard-MIT) for detailed discussion and additional data regarding the FlowSeq datasets. We would also like to thank Robert Egbert (Berkeley), Heather Beck (Vienna), Mark Mimee (MIT), Allison Hoynes-O’Connor (Washington U), and our lab members for discussion and contributions to the genetic system database.

## AUTHOR CONTRIBUTIONS

ACR and HMS developed the method, analyzed data, and wrote the manuscript. ACR compiled the database and developed the automated model test system software. HMS conceived the project and provided guidance on test system development.

## COMPETING FINANCIAL INTERESTS

ACR declares no competing financial interests. HMS is the founder of De Novo DNA.

## METHODS

### Compiling the genetic system database

To create the database, we compiled sequences and expression measurements from the supplemental data of several publications. We included detailed information for all sequences including the position of the start codon, bacteria host, experimental conditions, and characterization method and data. For the 1014IC dataset, mean fluorescence (or luminescence) levels, standard deviation across at least three replicates, and RT-qPCR measurements (when available) were included. For the 8848FS dataset, raw data includes binned read counts, RNA read counts, and DNA read counts for two replicates. The reconstructed relative fluorescence values (protein levels) and the mRNA levels were recalculated as described in the original paper^29^. Apparent translation rates for the FlowSeq datasets are calculated as the protein level divided by the mRNA level.

DNA sequences for the genetic systems were either provided in the supplementary information of the publications, acquired directly from the original authors, or recreated manually from information found in the publication^25^.5’-untranslated regions (UTRs) of the mRNAs were determined by identifying the most common transcription start site (TSS) of the promoter of the genetic system. Each TSS was either estimated by selected the first nucleotide following the annotated promoter, or in the case of any promoters characterized by the 8848FS dataset, the TSS is defined as the most frequent upstream position of transcription initiation as identified by RNA read counts. Start codons specified by the original publications are saved in the database.

Data was first stored in a pandas dataframe (<u>http://pandas.pydata.org/</u>) with null values for entries where information or characterization data was not provided. The supporting model test system code includes functions to allow easy manipulation of the dataframe, such as extracting and analyzing genetic systems that share common properties (see SynBioMTS/dbms.py).

### Filtering the FlowSeq datasets

We applied a combination of filters on the FlowSeq dataset to remove characterized sequences with low read counts or skewed binned read counts. Following the rules filters in Kosuri *et al*.^29^, we removed sequences with insufficient protein data, or if the reconstructed fluorescence level was above or below the estimated measurement range. The thresholds were defined as two-fold the minimum protein level (1357 reconstructed RFU) and 99% of the maximum protein level (204,060 reconstructed RFU). Characterized sequences that had fewer than 10 DNA read counts (both replicates), fewer than 50 DNA read counts (either replicate), or fewer than 20 RNA read counts (either replicate) were excluded. Lastly, promoters that had transcription start sites that started after the barcode sequence were excluded. These filters removed 3319 sequences (26.2%), resulting in the 8848FS dataset.

### Calculating total sequence entropy

We calculated the sequence diversity of a dataset (*H*_*seq*_) by determining the sum of the Shannon entropy across all nucleotide positions from the transcriptional start site to 35 nucleotides after the start codon (Equation 1). Considering that sequences have different length 5’ UTRs, we padded the 5’ ends of all sequences with “X” characters until all sequences have the same length. We then calculated Shannon entropies using a 5-letter alphabet with a maximum entropy, log_2_(5) = 2.32 bits, at each position.

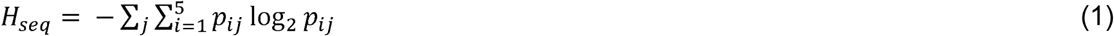

### Calculating Kullback-Leibler divergence

We calculated the model’s Kullback-Leibler (KL) divergence, which is the information gain achieved if the model of interest (*P*) is used instead of a random model (*Q*). We defined a random model (*Q*) as having a uniform error distribution across the observed range. We then calculate the relative probability of model error (*P*) across 100 log-spaced bins from 10^−3^ to 10^3^.

The *D*_*KL*_ value is computed using Equation (2), removing bins where *P(i)* is zero.The normalized *D*_*KL*_ value is divided by the maximum possible *D*_*KL*_ for a perfect model, which is 4.61 here.

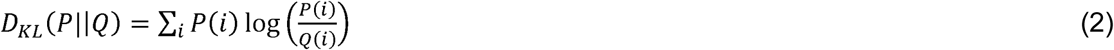

### Calculating area under the ROC curve (AUROC)

Measured protein expression levels are partitioned into HIGH and LOW binary outcomes according to a selected cutoff value. The test metric is the predicted translation initiation rate, using the same cutoff to determine LOW and HIGH predicted outcomes. To generate a Receiver Operating Characteristic curve, the cutoff is then systematically varied between the minimum and maximum of the measured protein expression levels, and the True Positive Rates (TPR) and False Positive Rates (FPR) are calculated for each cutoff value. The area under the ROC curve (AUROC) is then calculated using the trapezoidal rule (see calc_AUROC method).

### Using the gene expression models

All models were used with settings to maximize the predictive accuracy of each model, to best assess the proposed determinants. For all versions of RBS Calculator, the nine, 3’-most nucleotides of the 16S rRNA of each species was used as the aSD sequence, the temperature was set at the experimental temperature, a fixed mRNA post-cutoff of 35 nt was used, and all other parameters were left as default. To our knowledge, no source code is available for UTR Designer, so we recreated the model by modifying RBS Calculator v1.0 with the described changes^20^ (**Supplementary Discussion 1**) and ran the model with all the same parameters as RBS Calculator v1.0. For all versions of RBS Calculator and UTR Designer, we selected the most highly translated, in-frame start codon as the start site to predict the translation rate. RBS Designer was released as a Windows installer package. We wrote a Python wrapper to run the batch version of the program on the full database (see RBSBatch.py in the repository). All default settings were used and the correct rRNA sequences were used for RBS Designer. Noncanonical start codons (e.g. “CUG”) were skipped because RBS Designer only supports prediction for genes with an AUG start. For EMOPEC, we used the predicted Expression value for comparisons, rather than the Expression_percent_ value as described in Bonde *et al.*^25^. All default settings were used for EMOPEC. All model output was saved to the database for future study.

## References

1 Taylor, N. D. et al. Engineering an allosteric transcription factor to respond to new ligands.Nature methods 13, 177 (2016).

2 Espah Borujeni, A., Mishler, D. M., Wang, J., Huso, W. & Salis, H. M. Automated physics-based design of synthetic riboswitches from diverse RNA aptamers. Nucleic Acids Research 44, 1–13, doi:10.1093/nar/gkv1289 (2016).

3 Brophy, J. A. & Voigt, C. A. Principles of genetic circuit design. Nature methods 11, 508–520 (2014).

4 Kushwaha, M. & Salis, H. M. A portable expression resource for engineering cross-species genetic circuits and pathways. Nature communications 6 (2015).

5 Lee, S. Y. & Kim, H. U. Systems strategies for developing industrial microbial strains. Nature Biotechnology 33, 1061–1072, doi:10.1038/nbt.3365 (2015).

6 Smanski, M. J. et al. Synthetic biology to access and expand nature’s chemical diversity. Nature reviews. Microbiology 14, 135 (2016).

7 Ng, C. Y., Farasat, I., Maranas, C. D. & Salis, H. M. Rational design of a synthetic Entner– Doudoroff pathway for improved and controllable NADPH regeneration. Metabolic engineering 29, 86–96 (2015).

8 Brewster, R. C., Jones, D. L. & Phillips, R. Tuning Promoter Strength through RNA Polymerase Binding Site Design in Escherichia coli. PLoS Computational Biology 8, doi:10.1371/journal.pcbi.1002811 (2012).

9 Brewster, R. C. et al. The transcription factor titration effect dictates level of gene expression. Cell 156, 1312–1323 (2014).

10 Chen, Y.-J. et al. Characterization of 582 natural and synthetic terminators and quantification of their design constraints. Nature Methods 10, 659–663, doi:10.1038/nmeth.2515 (2013).

11 Espah Borujeni, A. & Salis, H. M. Translation initiation is controlled by RNA folding kinetics via a ribosome drafting mechanism. Journal of the American Chemical Society 138, 7016–7023 (2016).

12 Fernandez-Rodriguez, J., Moser, F., Song, M. & Voigt, C. A. Engineering RGB color vision into Escherichia coli. Nature Chemical Biology (2017).

13 Gupta, A., Reizman, I. M., Reisch, C. R. & Prather, K. L. Dynamic regulation of metabolic flux in engineered bacteria using a pathway-independent quorum-sensing circuit. Nat Biotechnol, doi:10.1038/nbt.3796 (2017).

14 Smanski, M. J. et al. Functional optimization of gene clusters by combinatorial design and assembly. Nature biotechnology 32, 1241–1249 (2014).

15 Nielsen, A. A. et al. Genetic circuit design automation. Science 352, aac7341 (2016).

16 Jacobson, I., Booch, G., Rumbaugh, J., Rumbaugh, J. & Booch, G. The unified software development process. Vol. 1 (Addison-wesley Reading, 1999).

17 Moult, J. A decade of CASP: progress, bottlenecks and prognosis in protein structure prediction. Current Opinion in Structural Biology 15, 285–289, doi:10.1016/j.sbi.2005.05.011 (2005).

18 Prill, R. J. et al. Towards a Rigorous Assessment of Systems Biology Models: The DREAM3 Challenges. PLoS ONE 5, doi:10.1371/journal.pone.0009202 (2010).

19 Meyer, P. et al. Industrial methodology for process verification in research (IMPROVER): toward systems biology verification. Bioinformatics 28, 1193–1201, doi:10.1093/bioinformatics/bts116 (2012).

20 Seo, S. W. et al. Predictive design of mRNA translation initiation region to control prokaryotic translation efficiency. Metabolic Engineering 15, 67–74, doi:10.1016/j.ymben.2012.10.006 (2013).

21 Salis, H. M., Mirsky, E. A. & Voigt, C. A. Automated design of synthetic ribosome binding sites to control protein expression. Nature Biotechnology 27, 946–950, doi:10.1038/nbt.1568 (2009).

22 Na, D., Lee, S. & Lee, D. Mathematical modeling of translation initiation for the estimation of its efficiency to computationally design mRNA sequences with desired expression levels in prokaryotes. BMC Systems Biology 4, 1–16, doi:10.1186/1752-0509-4-71 (2010).

23 Farasat, I. et al. Efficient search, mapping, and optimization of multi-protein genetic systems in diverse bacteria. Molecular Systems Biology 10, 1–18 (2014).

24 Espah Borujeni, A., Channarasappa, A. S. & Salis, H. M. Translation rate is controlled by coupled trade-offs between site accessibility, selective RNA unfolding and sliding at upstream standby sites. Nucleic Acids Research 42, 2646–2659, doi:10.1093/nar/gkt1139 (2014).

25 Bonde, M. T. et al. Predictable tuning of protein expression in bacteria. Nature Methods 13, 233–236, doi:10.1038/nmeth.3727 (2016).

26 Espah Borujeni, A. et al. Precise quantification of translation inhibition by mRNA structures that overlap with the ribosomal footprint in N-terminal coding sequences. Nucleic Acids Res, doi:10.1093/nar/gkx061 (2017).

27 Tian, T. & Salis, H. M. A predictive biophysical model of translational coupling to coordinate and control protein expression in bacterial operons. Nucleic Acids Research 43, 7137–7151, doi:10.1093/nar/gkv635 (2015).

28 Mimee, M., Tucker, A. C., Voigt, C. A. & Lu, T. K. Programming a Human Commensal Bacterium, Bacteroides thetaiotaomicron, to Sense and Respond to Stimuli in the Murine Gut Microbiota. Cell Systems 1, 62–71, doi:10.1016/j.cels.2015.06.001 (2015).

29 Kosuri, S. et al. Composability of regulatory sequences controlling transcription and translation in Escherichia coli. Proceedings of the National Academy of Sciences 110, 14024–14029 (2013).

30 Goodman, D. B., Church, G. & Kosuri, S. Causes and Effects of N-Terminal Codon Bias in Bacterial Genes. Science 342, 475–479 (2013).

31 Espah Borujeni, A. & Salis, H. M. Translation Initiation is Controlled by RNA Folding Kinetics via a Ribosome Drafting Mechanism. Journal of the American Chemical Society 138, 7016–7023, doi:10.1021/jacs.6b01453 (2016).

32 Beck, H. J., Fleming, I. M. & Janssen, G. R. 5’-Terminal AUGs in Escherichia coli mRNAs with Shine-Dalgarno Sequences: Identification and Analysis of Their Roles in Non-Canonical Translation Initiation. PloS one 11, e0160144 (2016).

33 Beck, H. J. & Janssen, G. R. Novel translation initiation regulation mechanism in Escherichia coli ptrB mediated by a 5’-terminal AUG. Journal of Bacteriology, JB. 00091–00017 (2017).

34 Egbert, R. G. & Klavins, E. Fine-tuning gene networks using simple sequence repeats. Proceedings of the National Academy of Sciences 109, 16817–16822 (2012).

35 Hecht, A. et al. Measurements of translation initiation from all 64 codons in E. coli. Nucleic acids research 45, 3615–3626 (2017).

36 Marzi, S. et al. Structured mRNAs regulate translation initiation by binding to the platform of the ribosome. Cell 130, 1019–1031, doi:10.1016/j.cell.2007.07.008 (2007).

37 Kozak, M. Regulation of translation via mRNA structure in prokaryotes and eukaryotes. Gene 361, 13–37 (2005).

38 Kozak, M. Initiation of translation in prokaryotes and eukaryotes. Gene 234, 187–208 (1999).

39 Vellanoweth, R. L. & Rabinowitz, J. C. The influence of ribosome-binding-site elements on translational efficiency in Bacillus subtilis and Escherichia coli in vivo. Molecular Microbiology 6, 1105–1114, doi:10.1111/j.1365-2958.1992.tb01548.x (1992).

40 Hongyun Chen, M. B., Ravindra Kumar and Ernest Jay. Determinantion of the optimal aligned spacing between the SD sequence and the start codon of E coli mRNAs. Nucleic Acids Res 22, 4953–4957 (1994).

41 Crooks, G. E., Hon, G., Chandonia, J.-M. & Brenner, S. E. WebLogo: A Sequence Logo Generator. Genome Research 14, 1188–1190, doi:10.1101/gr.849004.1 (2004).

42 Andronescu, M., Condon, A., Hoos, H. H., Mathews, D. H. & Murphy, K. P. Computational approaches for RNA energy parameter estimation. RNA 16, 2304–2318 (2010).

43 Sprengart, M. L., Fatscher, H. P. & Fuchs, E. The initiation of translation in E. coli: apparent base pairing between the 16srRNA and downstream sequences of the mRNA. Nucleic Acids Research 18, 1719–1723 (1990).

44 Wegmann, U., Horn, N. & Carding, S. R. Defining the Bacteroides Ribosomal Binding Site. Applied and Environmental Microbiology 79, 1980–1989, doi:10.1128/AEM.03086-12 (2013).

45 Dykeman, E. C. An implementation of the Gillespie algorithm for RNA kinetics with logarithmic time update. Nucleic Acids Research 43, 5708–5715, doi:10.1093/nar/gkv480 (2015).

46 Mackie, G. A. RNase E: at the interface of bacterial RNA processing and decay. Nature Reviews Microbiology 11, 45–57, doi:10.1038/nrmicro2930 (2013).

47 Deana, A. & Belasco, J. G. Lost in translation: the influence of ribosomes on bacterial mRNA decay. Genes & development 19, 2526–2533 (2005).

48 Espah Borujeni, A. The development of equilibrium and non-equilibrium models of translation and riboswitch regulation: Towards the automated design of cellular sensors, The Pennsylvania State University, (2015).

